# FrozenChicken: Promoting the meta-analysis of chicken microarray data

**DOI:** 10.1101/2021.02.25.432894

**Authors:** Isabel Duarte, Marta Liber, Ramiro Magno, Raquel P. Andrade

## Abstract

The FrozenChicken RData package, contains the frozen vectors for the commercially available (in situ oligonucleotide) Affymetrix Chicken Genome Array (GEO platform id GPL3213). This package will promote, simplify, and ease the meta-analysis of chicken microarray data by the research community studying vertebrate development using the chick model organism. The package is freely available in https://github.com/iduarte/FrozenChicken. (\**Equal contribution*.)

## Introduction

### Background

*Gallus gallus* (chicken) is one of the most valuable model organisms for the study of the vertebrate embryo development. Such studies can be aided by pooling together OMICs data from public repositories, like GEO (Gene Expression Omnibus) and ArrayExpress, that currently contain more than 11.660 datasets from chicken, representing a wealth of data that can be explored to answer fundamental questions and generate new hypotheses. However, since these data come from different experiments, their meta-analysis requires proper normalization to deal with the technical biases and batch effects before making the data comparable for statistical analysis.

### Approach

An effective method for such normalization is the single-array pre-processing provided by the Frozen Robust Multiarray Analysis (**fRMA**) (McCall MN, et al. Biostatistics. 2010). This method uses *“frozen” RMA* vectors pre-computed from great amounts of available data for the same microarray platform, accounting for the aforementioned biases. However, such frozen vectors/parameters are available for multiple organisms (most notably, human, mouse, zebrafish, and fruit-fly among others), but not for chicken, hence preventing, or delaying, the proper meta-analysis of chicken transcriptomics datasets without prior computation of self-generated frozen parameters.

### Output

Here, we present the RData package **FrozenChicken** containing the chicken microarray frozen vectors that can be directly plugged-in to a chicken microarray analysis pipeline (using fRMA) without any other prior data gathering and processing. The package is freely available for the research community at: https://github.com/iduarte/FrozenChicken. A version of this article in the form of an html tutorial has been deposited in zenodo DOI:10.5281/zenodo.3765944.

### Significance

This package will directly benefit the chicken research community by facilitating future meta-analysis studies using transcriptomics datasets from public repositories, hence directly contributing to the quality of the scientific research using the chick model organism.

## Methods

### Methods | 1. Data Collection

#### 1 Search for relevant GEO data series

The chicken microarray datasets used were gathered using the **ESearch** function from the **Entrez** Programming Utilities (E-utilities) that provide a programmatic connection with the Entrez query system from NCBI.

This search returned a list of Unique Identifiers (UIDs) for **1739 records** that met the following query criteria:

- search the database Geo DataSets (*gds*);
- search for GEO platform id GPL3213, which is the **Affymetrix Chicken Genome Array** (the chicken commercially available microarray chip);
- select only records that have .*CEL* supplementary files available for downloading.

The issued query was the following: https://eutils.ncbi.nlm.nih.gov/entrez/eutils/esearch.fcgi?db=gds&term=GPL3213%5BACCN%5D+AND+cel%5BsuppFile%5D&retmax=5000&usehistory=y

#### 2 Gather summary data for the data series found

Using the list of UIDs returned from the previous step, we gathered the metadata associated with each entry. For this we used the **ESummary** function from NCBI’s **E-utilities**, that returns the documented summaries for each UID, including the Geo Series (GSE) identifier for each experiment. The ESummary tool directly uses the esearch URL retrieved from the previous step.

#### 3 Download the relevant data sets

Careful manual inspection of the results retrieved from the previous step, led to the collection of **128 relevant GSE records**. These were downloaded using the R package GEOquery (version 2.50.5) (https://academic.oup.com/bioinformatics/article/23/14/1846/190290).

~~~
## Load the required package GEOquery
library (GEOquery)
## Set the working directory to the folder where the downlods will be
saved setwd(“/home/FrozenChicken/data”)
## Create the vector of GSE ids to download
GSE.ids <- c(“GSE94622”,”GSE114476”,”GSE89325”,”GSE106788”,”GSE106787”,
        “GSE109451”,”GSE81023”,”GSE87663”,”GSE79963”,”GSE69862”,
        “GSE71888”,”GSE87486”,”GSE51330”,”GSE76794”,”GSE82344”,
        “GSE48454”,”GSE48453”,”GSE69684”,”GSE81994”,”GSE39450”,
        “GSE81717”,”GSE81461”,”GSE75798”,”GSE35430”,”GSE14220”,
        “GSE71117”,”GSE62882”,”GSE53932”,”GSE53931”,”GSE53930”,
        “GSE33389”,”GSE60754”,”GSE31507”,”GSE59002”,”GSE59921”,
        “GSE59920”,”GSE48359”,”GSE48116”,”GSE52227”,”GSE34687”,
        “GSE14587”,”GSE44394”,”GSE50880”,”GSE22222”,”GSE47191”,
        “GSE38168”,”GSE31508”,”GSE31524”,”GSE31506”,”GSE31505”,
        “GSE31501”,”GSE31499”,”GSE31476”,”GSE37070”,”GSE42845”,
        “GSE39602”,”GSE42516”,”GSE40802”,”GSE40100”,”GSE398242”,
        “GSE39346”,”GSE27958”,”GSE35581”,”GSE38381”,”GSE38107”,
        “GSE37782”,”GSE35413”,”GSE15830”,”GSE32272”,”GSE21706”,
        “GSE25185”,”GSE25151”,”GSE32494”,”GSE24641”,”GSE25588”,
        “GSE29565”,”GSE29564”,”GSE29563”,”GSE29562”,”GSE28634”,
        “GSE28391”,”GSE28388”,”GSE21915”,”GSE22592”,”GSE23592”,
        “GSE23881”,”GSE23389”,”GSE19698”,”GSE22230”,”GSE17758”,
        “GSE17725”,”GSE21679”,”GSE15143”,”GSE15141”,”GSE14489”,
        “GSE14013”,”GSE14509”,”GSE11636”,”GSE18477”,”GSE18778”,
        “GSE18568”,”GSE18506”,”GSE126752”,”GSE16081”,”GSE16064”,
        “GSE15413”,”GSE15382”,”GSE9251”,”GSE11597”,”GSE10538”,
        “GSE12268”,”GSE11439”,”GSE8010”,”GSE8483”,”GSE8018”,
        “GSE8017”,”GSE8016”,”GSE10231”,”GSE8495”,”GSE9884”,
        “GSE8693”,”GSE7805”,”GSE6543”,”GSE7176”,”GSE6856”,
        “GSE6844”,”GSE6843”,”GSE6868”)
## Run getGEOSuppFiles function from the GEOquery function
sapply(GSE.ids, getGEOSuppFiles)
~~~

### Methods | 2. Building the FrozenChicken Package

To create the FrozenChicken R package we used the frmaTools R package (version 1.34.0) (https://doi.org/10.1186/1471-2105-12-369). This package requires the usage of the same number of samples per experiment in order not to create biases. We chose to use **4 samples** from each experiment, keeping only the datasets that had at least four microarrays.

Therefore, we used **118 Geo DataSets** from Affymetrix Chicken Genome Array platform, from which 4 arrays, per batch, were randomly selected (**Batch number = 118** and **Batch size = 4**). This approach retrieved a powerful set of **472 chips** that were used as input for the frmaTools R package to compute the frozen vectors/parameters.

~~~
## Load the required packages
library (frmaTools)
library (chickencdf) # Annotation package for the chicken microarray
## Set the directories required
frma_data_dir <- “/home/FrozenChicken/data”
frma_output_dir <- “/home/FrozenChicken/output”
## Create a numerical vector for the batch size
frma_batch_size <- 4
## Create the batch-id string
fRMA.chiken.batch.id <- rep(1:(length(dir(frma_data_dir))/frma_batch_size),
                                  each=frma_batch_size)
## Run the makeVectorPackage function from the frmaTools package
makeVectorPackage (dir(frma_data_dir), fRMA.chiken.batch.id,
                   file.dir=frma_data_dir, output.dir=frma_output_dir,
                   version=“1.0”, maintainer=“Marta Liber <mliber.pt@gmail.com>“,
                   species=“Gallus gallus”, annotation=“chickencdf”,
                   packageName=“affyChickGenomeArrayfrmavecs”, background=“rma”,
                   normalize=“quantile”, normVec=NULL, type=“AffyBatch”,
                   unlink=TRUE, verbose=TRUE)
~~~

## Results

### Case Study Using FrozenChicken

In this section we will show how to use the FrozenChicken vectors in a microarray data analysis of chicken transcriptomics. The workflow described here was performed in R, using RStudio (version 1.1.463). The typical workflow of a microarray data analysis is shown in Figure 1.

**Figure 1:**
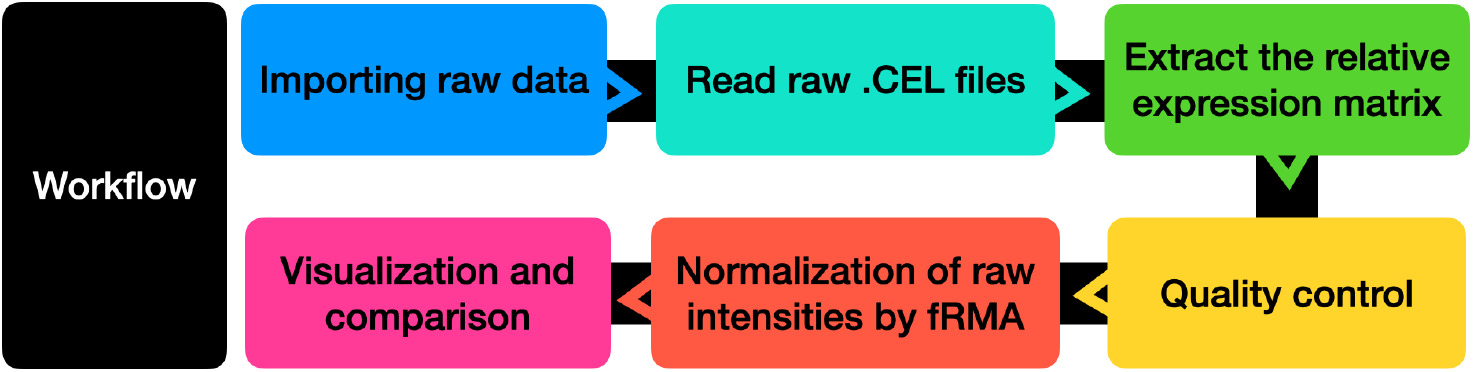
Microarray data analysis typical workflow.

**Figure 2:**
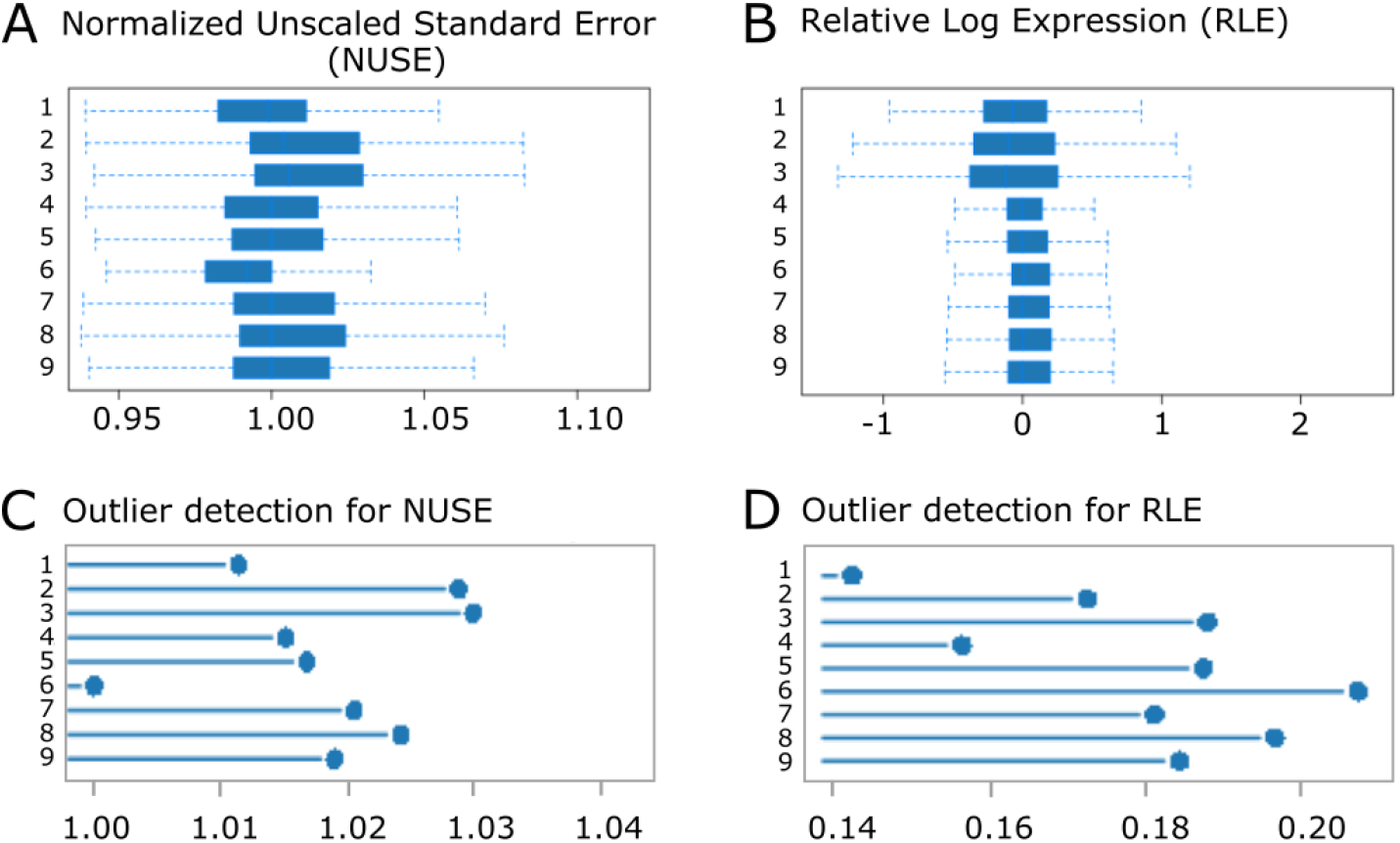
Quality Control metrics report. (A) Normalized Unscaled Standard Error (NUSE) plot. (B) Relative Log Expression (RLE) plot. (C) Bar chart representation for the outlier detection based on NUSE metrics in (A). (D) Bar chart representation for the outlier detection based on RLE metrics in (B).

Here, we will be looking at three chicken microarray datasets from different experiments:

- chick PSM tissue, embryo stage HH12, from GEO record GSE75798 (Oginuma, M. *et al*., 2017);
- chick right PSM region, embryo stages HH11-12, from Array Express record E-MTAB-406 (Krol *et al*., 2011);
- chick anterior and posterior limb bud at stage HH20, from ArrayExpress record E-MTAB-4048 (Anderson, C. *et al*., 2016).

#### Case study | 1. Install FrozenChicken and Additional R Packages

We will start by installing the FrozenChicken package, which is deposited in GitHub. To install it directly from GitHub you should use the package remotes. If you do not have it, install it first:

~~~
## Install the package from CRAN repository
install.packages(“remotes”)
## Load the package
library(remotes)
~~~

Then install the R package FrozenChicken directly from GitHub:

~~~
remotes::install_github(“iduarte/FrozenChicken”)
~~~

Next you can load the library named **affyChickGenomeArrayfrmavecs** and the *frozen parameters* become available for the normalization of chicken microarray data from different experiments (provided that all use the same Affymetrix Chicken Genome Array platform).

~~~
## Load the FrozenChicken package
  # This is the full name of the FrozenChicken data object
library(affyChickGenomeArrayfrmavecs)
## Load the affyChickGenomeArrayfrmavecs data set
data(affyChickGenomeArrayfrmavecs)
~~~

To complete this case study, the following R Packages are required:

~~~
## Install Bioconductor (if not already installed)
if (!requireNamespace(“BiocManager”, quietly = TRUE))
    install.packages(“BiocManager”)
BiocManager::install(version = “3.10”)
## Install the required Packages (if not already installed)
BiocManager::install(c(“ArrayExpress”,
                              “GEOquery”,
                              “Biobase”,
                              “affy”,
                              “arrayQualityMetrics”,
                              “ggplot2”,
                              “frma”,
                              “devtools”))
~~~

#### Case Study | 2. Obtaining the Gene Expression Matrix

To conduct a transcriptomics data analysis, one must obtain a gene expression matrix, i.e. a data table that reports the expression level measured for each gene. When the data to be analysed originates from microarrays that are deposited in public repositories, namely GEO or ArrayExpress, the completion of the following steps will generate a gene expression matrix:

1. Download the .CEL files (raw microarray data from Affymetrix) from its data repository. The data can be download using the ArrayExpress and GEOquery packages, respectively.
2. Read the raw .CEL files into R using the affy package, creating an ‘AffyBatch’ object containing the microarray data.
3. Extract the gene expression matrix from the ‘AffyBatch’ object.

~~~
## Load the required packages for this code chunk
library(devtools)
library(ArrayExpress)
library(GEOquery)
library(Biobase)
library(affy)
## Set up the directories used for the analysis
  ## (NOTE: Change the paths to the correct directories from your computer)
setwd (“/home/microarray_meta_analysis/”)
data_dir <- “/home/microarray_meta_analysis/data”
output_dir <- “/home/microarray_meta_analysis/output”
## Step 1 - Downloading the .CEL files
setwd (data_dir)
getGEOSuppFiles (“GSE75798”, makeDirectory = TRUE)
getAE (“E-MTAB-4048”, type = “raw”)
getAE (“E-MTAB-406”, type = “raw”)
## Step 2 - Load the .CEL files, i.e. import the raw data into R
affybatch_chick <- ReadAffy (celfile.path = data_dir)
## Step 3 - Extract the raw expression values using the exprs() function
expres_chick_raw <- exprs (affybatch_chick)
# Log2 transform (only if values are not log already)
  # NOTE: Code adapted from NCBI’s GEO2R scripts.
qx <- as.numeric(quantile(expres_chick_raw,
                                 c(0., 0.25, 0.5, 0.75, 0.99, 1.0), na.rm=T))
LogC <- (qx[5] > 100) ||
  (qx[6]-qx[1] > 50 && qx[2] > 0) ||
  (qx[2] > 0 && qx[2] < 1 && qx[4] > 1 && qx[4] < 2)
# log2 transform values if they are not in log scale already
if (LogC) {
 expres_chick_raw[which(expres_chick_raw <= 0)] <- NaN # remove zeros
 expres_chick_raw <- log2(expres_chick_raw)
 cat(“The RAW expression values were not log2 transformed,”,
             “and now they have been log2 transformed.”)
}
# Cleanup (delete unnecessary variables)
rm (qx, LogC)
~~~

#### Case Study | 3. Quality Control

Quality control (QC) is an important step to remove data from faulty arrays. The arrayQualityMetrics package flags potential outliers and outputs plots to aid with the visual inspection of the results.

All arrays passed the QC criteria, and so all will be included in the next analysis steps. If any outlier were to be flagged, then those arrays should be removed from the analysis, and the quality control steps have to be re-run.

~~~
## Load the required library for quality control
library(arrayQualityMetrics)
## Step 3. Quality Control Report
  # This package uses the raw affybatch object directly
  # and not the expression matrix
  # (which is why we log transform the data).
arrayQualityMetrics(expressionset = affybatch_chick,
                    outdir = output_dir,
                    force = FALSE, do.logtransform = TRUE)
~~~

#### Case Study | 4. Normalization with fRMA Using FrozenChicken

The normalization of data obtained from different experiments is pivotal to make the data comparable between arrays. Using the frozenRMA method this can be easily done using a vector of frozen parameters pre-computed from diverse datasets from the same microarray chip. FrozenChicken presents a package containing the pre-computation of these frozen parameters for the chicken commercial microarray **Affymetrix Chicken Genome Array** to be used with the `fRMA` package.

~~~
## Load the required packages
library(frma)
library(affyChickGenomeArrayfrmavecs)
## Load the affyChickGenomeArrayfrmavecs data set to use latter
data(affyChickGenomeArrayfrmavecs)
## Step 4 - frozenRMA normalization using the FrozenChicken vectors
eset_chick_frma <- frma(affybatch_chick,
                               background=“rma”,
                               normalize=“quantile”,
                               summarize=“robust_weighted_average”,
                               target=“probeset”,
                               input.vecs=affyChickGenomeArrayfrmavecs,
                               output.param=NULL, verbose=FALSE)
# The data is Log2 tranformed by the process of fRMA normalization
expres_chick_frma <- exprs(eset_chick_frma)
~~~

#### Case Study | 5. Data Visualization

Once the data have been normalized, we must confirm that the normalization was successful by running the quality control steps on the newly normalized data, and compare the results with the pre-normalized data. Here we show two of the most relevant plots to evaluate the success of the normalization procedure, namely, a boxplot (where each box corresponds to the intensity distribution of one array), and a Principal Component Analysis PCA plot to view the variation between the arrays (here, each dot is one array).

~~~
## Load the required package for this code chunk
library(ggplot2)
# Boxplots of raw log-intensity distribution
# requires the raw expression values extracted in step 2
boxplot(expres_chick_raw, col=c(rep(“#b2df8a”,3),
                                rep(“#1f78b4”,3),
                                rep(“#fb9a99”,3)),
       las = 3, cex.axis=0.75,
       main=“Chicken Log2 raw expression values”)
## PCA
pca_chicken <- prcomp(t(expres_chick_raw),
                             center = TRUE,
                             scale. = TRUE)
pca_chicken_information <- data.frame(pca_chicken$x,
                             variance=as.numeric(round(
                              100*summary(pca_chicken)$importance[2,],
                              digits=2)),
                       origin=c(rep(“PSM AE”, 3),
                                rep(“PSM GEO”,3),
                                rep(“Limb AE”,3)))
ggplot(pca_chicken_information,
       aes(x=pca_chicken_information[,1],
           y=pca_chicken_information[,2],
           color=origin)) +
 geom_point(size=2, alpha=0.7, show.legend = TRUE) + theme_bw() +
 labs(color=‘Origin’) +
 scale_color_manual(values = c(“#b2df8a”, “#1f78b4”, “#fb9a99”)) +
 xlab(paste(“PC1 (“, pca_chicken_information$variance[1], “%)”)) +
 ylab(paste(“PC2 (“, pca_chicken_information$variance[2], “%)”)) +
 ggtitle(“Chicken Log2 raw expression values”) + geom_hline(yintercept = 0) +
 geom_vline(xintercept = 0) -> pca_plot_chicken
pca_plot_chicken
## Boxplots of normalized log-intensity distribution
# requires the normalized expression values extracted in step 4
boxplot(expres_chick_frma,
        col=c(rep(“#b2df8a”,3),
              rep(“#1f78b4”,3),
              rep(“#fb9a99”,3)),
        las = 3, cex.axis=0.75,
        main=“Chicken Log2 normalized expression values”)
        # PCA: provides another view of the correlations of expression between arrays.
        pca_chicken_norm <- prcomp(t(expres_chick_frma),
center = TRUE,
scale. = TRUE)
pca_chicken_norm_information <- data.frame(pca_chicken_norm$x,
                                                  variance=as.numeric(round(
                                                   100*summary(pca_chicken_norm)$importance[2,],
                                                   digits=2)),
                                                  origin=c(rep(“PSM AE”, 3),
                                                           rep(“PSM GEO”,3),
                                                           rep(“Limb AE”,3)))
ggplot(pca_chicken_norm_information,
       aes(x=pca_chicken_norm_information[,1],
           y=pca_chicken_norm_information[,2],
           color=origin)) +
 geom_point(size=2, alpha=0.7, show.legend = TRUE) +
 theme_bw() +
 labs(color=‘Origin’) +
 scale_color_manual(values=c(“#b2df8a”, “#1f78b4”, “#fb9a99”)) +
 xlab(paste(“PC1 (“, pca_chicken_norm_information$variance[1], “%)”)) +
 ylab(paste(“PC2 (“, pca_chicken_norm_information$variance[2], “%)”)) +
 ggtitle(“Chicken Log2 normalized expression values”) +
 geom_hline(yintercept = 0) +
 geom_vline(xintercept = 0) -> pca_plot_chicken_norm
 pca_plot_chicken_norm
~~~

At this point, the data is log2-transformed and normalized, therefore, further analyses on the transcriptomics dataset may be performed (e.g. differential gene expression, and functional enrichment).

### FrozenChicken Performance Evaluation

Before normalization, samples show variation between and within batches (Figure 3A). Additionally, PCA analysis found that 92.64% of variance between the data points is explained by the identity of the experiment (Figure 3C). Thus, the major source of variation in the raw intensity measurements is due to batch effects that should be reduced after the fRMA normalization.

**Figure 3:**
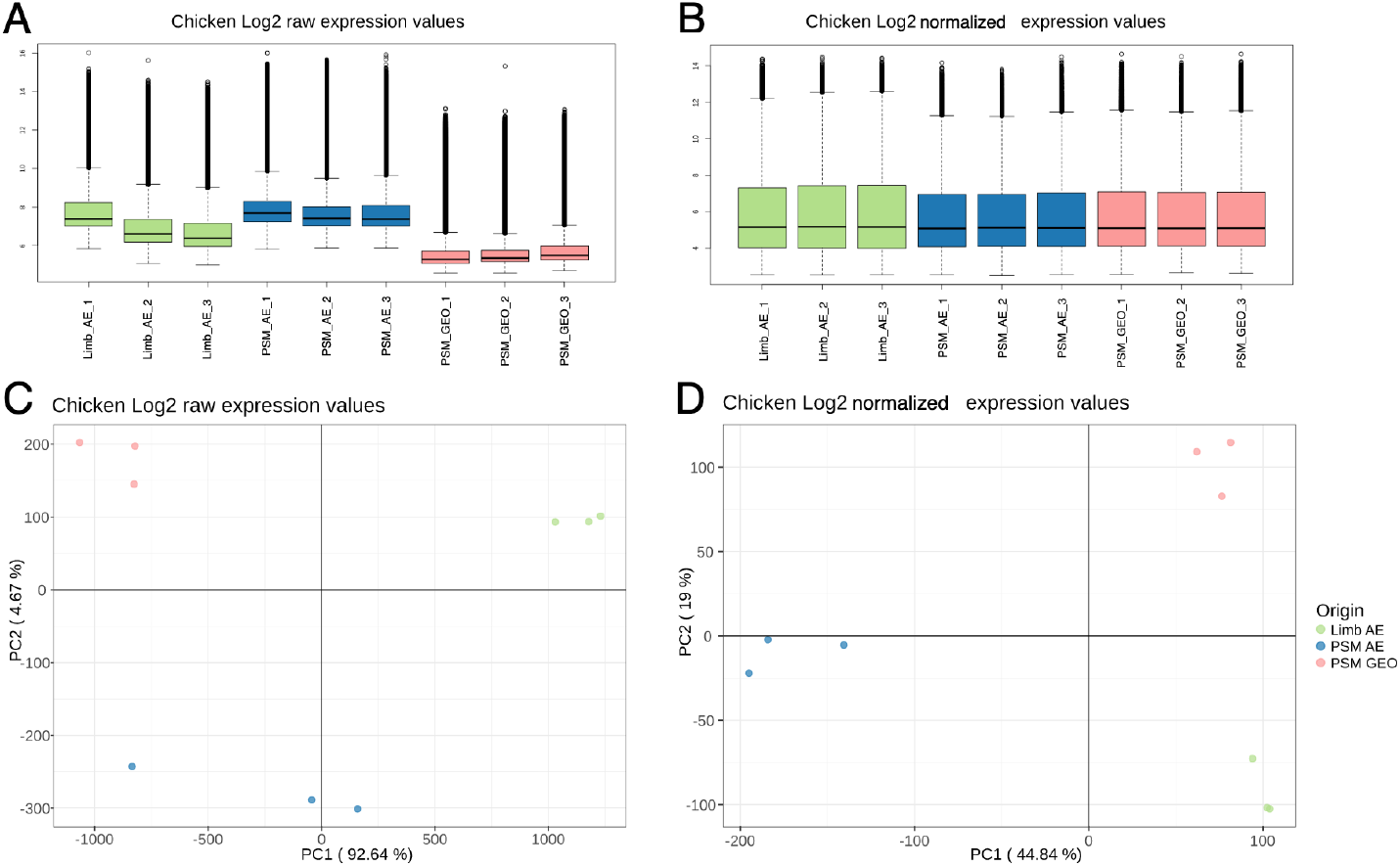
Comparison between raw and normalized gene expression values. (A) Boxplot of raw log2-transformed expression values. (B) Boxplot of normalized log2-transformed expression values. (C) PCA of raw log2-transformed expression values. (D) PCA of normalized log2-transformed expression values.

Since the purpose of normalization is to remove unwanted variation between the transcriptional profiles, we expect that after the normalization, the relative gene-expression estimates will be distributed in a homogeneous way across the arrays, and also, the variance found by the PCA will decrease.

Our results show that, after normalizing the samples with the frozen vectors from FrozenChicken, the arrays exhibit similar distribution profiles (Figure 3B), indicating that the normalization was successful.

In the PCA analysis, as expected, the variance described by the first component (PC1) has now decreased to 44.84% (Figure 3D). Additionally, the distances between the points in the first principal component from the raw data, range between −1500 and 1500 (Figure 3C), while the distances for the normalized values has decreased by nearly 10 fold (ranging between −200 and 100 (Figure 3D), further confirming the success of the fRMA normalization using the pre-computed parameters from FrozenChicken.

It should be noted that, despite the successful normalization, there are still variation in the data (Figure 3D), mostly explained by the difference in tissue types (Figure 3D and Figure 4B), i.e. the biological variability that we are interested in studying. In the PCA from the raw data, the samples cluster by data repository, showing that the major source of variation explained by the first component was the experiment (technical variation that we are not interested in studying) (Figure 3C and Figure 4A).

**Figure 4:**
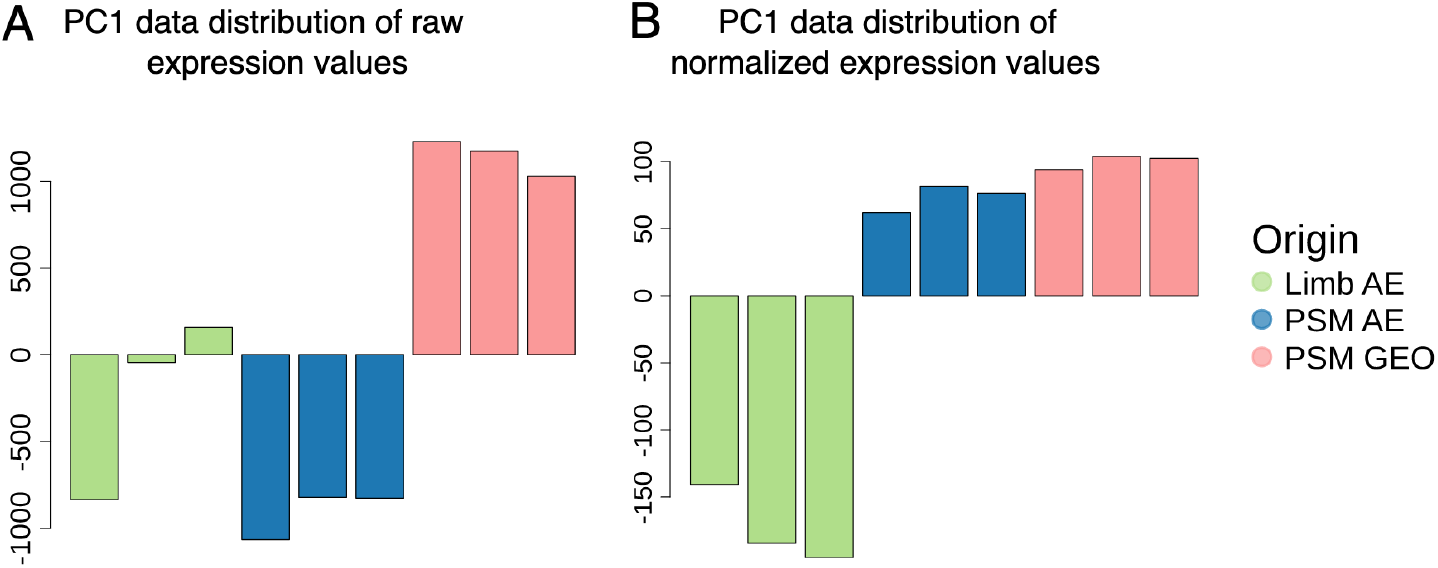
Distribution of distances calculated by PCA along its first component. (A) Distribution of distances along the first component of PCA on raw expression values. (B) Distribution of distances along the first component of PCA on normalized expression values.

## Conclusion

This case study shows that FrozenChicken is a **reliable data package** to be used with fRMA normalization for the pre-processing steps of chicken microarray data from different experiments, therefore promoting, simplifying, and easing future meta-analyses of chicken transcriptomics datasets from public repositories. This package will specially benefit the chicken research community, directly contributing to the quality of the scientific research using the chicken model organism. At the time of this publication (February 2021), the zenodo tutorial (DOI:10.5281/zenodo.3765944) describing this package had been downloaded over 1820 times (in less than one year), showing that our package has attracted the attention of our target audience.

## Author Contributions

ML collected the data, conducted the analysis, and wrote the manuscript. RM supervised the analysis, provided technical help, and wrote the manuscript. RPA devised the idea for the project, helped with interpreting the results, and wrote the manuscript. ID designed and supervised the analysis, interpreted the results, and wrote the paper. **\*ID and ML contributed equally to this work (ordered alphabetically)**. All authors read and approved the final manuscript.

## Acknowledgements

The authors would like to thank the members of the Temporal Control of Cell Differentiation Lab for precious feedback and insightful scientific discussions.

